# Coatomer protein complex I is required for efficient secretion of dengue virus non-structural protein 1

**DOI:** 10.1101/2024.10.12.618008

**Authors:** Stephen M. Johnson, Siena M. Centofanti, Gustavo Bracho, Michael R. Beard, Jillian M. Carr, Nicholas S. Eyre

## Abstract

Secreted non-structural protein 1 (sNS1) is an important orthoflavivirus pathogenic factor that can induce vascular leakage; a key symptom of severe dengue disease. Given the role of sNS1 in dengue pathogenesis, defining the molecular mechanisms of NS1 secretion may contribute towards development of NS1-targeting antiviral therapies. To this end, we performed a customised membrane-trafficking siRNA screen to identify human host factors involved in NS1 secretion. Our screen identified COPA, COPB2, and COPG1 as the top-ranking hits. These proteins are three of the seven subunits of the coatomer protein complex I (COPI) that coat transport vesicles that operate within the early secretory pathway, implicating COPI machinery as being involved in NS1 secretion. Validation studies employing host gene knockdown in dengue virus (DENV)-infected cells confirmed that COPI components are required for efficient NS1 secretion but are dispensable for infectious virus egress. Similar reductions in NS1 secretion were observed when COPI components were depleted in cells infected with West Nile virus Kunjin subtype (WNV/KUNV), indicating that the molecular mechanisms exploited to achieve NS1 secretion may be a conserved feature within the Orthoflavivirus genus. Heterologous expression of wildtype and pathogenic COPI variants in DENV NS1-NS5 polyprotein expressing cells resulted in altered NS1 secretion profiles, suggesting that allelic variants and altered expression levels of COPI components may indirectly influence the severity of dengue disease. The identification of COPI components as important determinants of NS1 secretion efficiency may aid in the identification of novel targets for anti-orthoflaviviral therapies.

**IMPORTANCE:** Over half of the world’s population is at risk of infection with mosquito-borne pathogenic orthoflaviviruses such as DENV. Although the secreted form of the viral NS1 protein has been identified as a major determinant of the pathogenic effects of DENV and related orthoflaviviruses, the exact mechanisms involved in NS1 secretion are poorly understood. Here we interrogated host factors involved in secretion of NS1 from infected cells using a customised membrane trafficking siRNA screen. This revealed 3 components of the COPI complex that regulates vesicular transport in the early secretory pathway as important factors in NS1 secretion. The involvement of COPI components in NS1 secretion was further validated using wildtype DENV and WNV/KUNV infection, overexpression approaches and chemical inhibition studies. Together, this study demonstrates the importance of COPI machinery in NS1 secretion and suggests that exploitation of this machinery in NS1 secretion may represent a future target of antiviral drug development.

## INTRODUCTION

Dengue virus (DENV) is the most prevalent arthropod-borne human viral pathogen, with half of the world’s population living in at-risk areas (1). It has been estimated that approximately 390 million DENV infections occur annually, with approximately 96 million infections resulting in disease (2). Symptoms range from mild febrile illness to life-threatening complications including dengue fever (DF), dengue haemorrhagic fever (DHF) and dengue shock syndrome (DSS). No dengue-specific antivirals are currently available.

DENV is an enveloped positive-sense single stranded RNA virus with an approximately 11 kb genome that encodes three structural proteins (Capsid, precursor-Membrane and Envelope) and seven non-structural proteins (NS1, NS2A, NS2B, NS3, NS4A, NS4B and NS5). Amongst these proteins, NS1 has attracted significant attention given its critical roles in viral RNA replication, infectious virus production and disease pathogenesis (3–5). Translated in the endoplasmic reticulum (ER), the hydrophilic NS1 monomer has two N-linked glycosylation sites, N130 and N207, and exists briefly as a soluble monomer. NS1 monomers rapidly dimerise forming a membrane-associated NS1 dimer, the predominant intracellular form (6, 7). Intracellular NS1 (iNS1) co-localises with dsRNA at both the ER-lumenal surface and interior of the virus-induced replication organelles where iNS1 plays an essential role in viral RNA replication (3, 8, 9). Recent evidence has demonstrated a role of iNS1 in virus particle assembly (10). NS1 dimers can also trimerize to form an open barrel-shaped NS1 hexamer that is stabilised by a lipid-rich central cavity and is efficiently secreted from the cell (11–13). Recent structural and biochemical studies, however, have revisited whether the extracellular sNS1 is hexameric, with studies reporting that sNS1 exhibits multiple oligomeric states including tetramers and hexamers (14, 15) and as dimers in complex with host serum factors(16, 17). It is this secreted form of NS1 that has garnered much attention recently for its role as a virulence factor being implicated in dengue disease pathogenesis (4, 18, 19). Plasma sNS1 levels, which correlate with viraemia in symptomatic individuals, have been shown to be significantly higher in patients who develop DHF than DF (20). Antibodies elicited against NS1 have been shown to contribute both protective and pathogenic consequences (21), and sNS1 acts in immune evasion by interfering with complement pathways (22, 23). Importantly, sNS1 is able to induce the release of vasoactive cytokines from immune cells and can directly induce endothelial cell glycocalyx layer disruption and vascular leakage (24, 25), a key symptom of severe dengue. Given the pathological effects of sNS1, much research has been conducted on the synthesis, structure, and key functional residues of this multifunctional protein (26, 27). However, major gaps exist in our understanding of the mechanisms that drive NS1 secretion.

To identify human host cell factors that are involved in NS1 secretion, we performed a customised membrane-trafficking siRNA screen. Our screen identified components of the coatomer protein complex I (COPI), a cage-like protein complex that coats transport vesicles that are best known for the bi-directional trafficking of proteins and lipids within the early secretory pathway (28–31). Validation studies confirmed that COPI is required for efficient NS1 secretion, and that the exploitation of these components to achieve NS1 secretion may be a conserved feature within the Orthoflavivirus genus. Further, we show that allelic variants of COPI may influence NS1 secretion profiles. This study expands our understanding of the molecular mechanisms of NS1 secretion and may aid in the identification of novel targets for anti-orthoflaviviral therapies.

## RESULTS

### A customised membrane-trafficking siRNA screen implicates COPI components as important determinants of DENV NS1 secretion

sNS1 is an important DENV pathogenic factor. Defining the molecular mechanisms of NS1 secretion may contribute towards the development of NS1-targeting antiviral therapies. To identify human host cell factors involved in DENV NS1 secretion we utilised an infectious DENV2-NS1-NLuc reporter virus construct that bears the small and sensitive NanoLuc luciferase (NLuc) reporter embedded within NS1 (9) (Fig. 1A). This reporter virus allows for the simple and reliable quantification of intracellular and secreted NS1-associated NLuc activity in infected cell cultures and is amenable to high-throughput screening.

**FIG 1.**
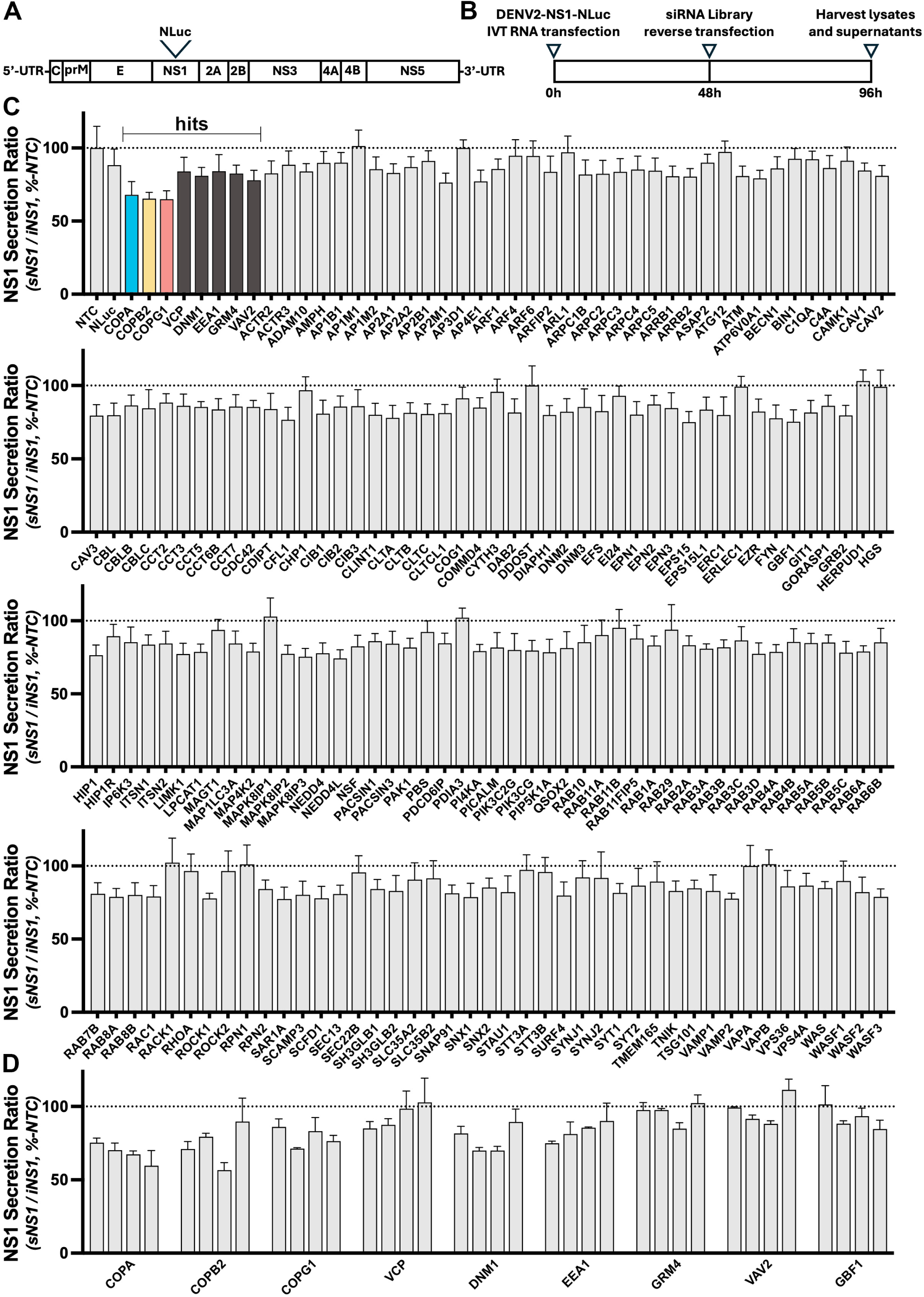
A customised membrane-trafficking siRNA screen implicates COPI components as important determinants of NS1 secretion. (A) Schematic overview of the infectious DENV2 construct bearing a NanoLuc luciferase (NLuc) tag within NS1. (B) Schematic overview of the high-throughput siRNA screen strategy. (C – D) Secretion ratio of NS1-NLuc (supernatant NS1-NLuc / lysate NS1-NLuc) as a % of average values associated with a non-targeting control (NTC) siRNA for the customised membrane-trafficking screen involving siRNA pools for each gene of interest (C) and deconvolution screen involving testing of the 4 individual siRNA duplexes that comprised the siRNA pools for hits identified in the primary screen, compared to the average values of those of the NTC control siRNA pool (D). For the membrane-trafficking siRNA screen (C), of each experimental siRNA pool, data are means + S.D. from nine measurements from three independent experiments. For the deconvolution screen (D), each experimental siRNA duplex that constituted the respective pool, data are means + S.D. from nine measurements from one experiment. Hit criteria for these siRNA screens are described in Supplementary Material.

The DENV2-NS1-NLuc reporter virus was used to screen a membrane-trafficking siRNA library (Dharmacon; 140 genes) that was customised and curated to include additional siRNA pools for 37 host genes which have been identified in previous studies as important host-dependency factors that may be manipulated by NS1 (32, 33). Each siRNA pool targeting the ∼180 host genes consists of four individual siRNA duplexes that recognise different sequences within each target transcript. Figure 1B provides a schematic overview of the siRNA screen strategy. Briefly, Firefly luciferase (FLuc)-expressing Huh-7.5 cells were transfected with *in vitro* transcribed DENV2-NS1-NLuc RNA and cultured for 48 hours to establish infection. Cells were then trypsinised and reverse transfected with the individual siRNA library pools in 96-well plates and returned to culture for a further 48 hours. Cell culture lysates and supernatants were harvested and assayed for intracellular and secreted NS1-associated NLuc activity, respectively, and normalised to intracellular Firefly luciferase activity, which was used as an indicator of cell viability. The data analysis methods and hit selection criteria are detailed in the Supplementary Material.

Only one experimental siRNA pool, RHOA, reproducibly reduced cell viability-associated FLuc luciferase levels to 1 standard deviation below the mean of the NTC. As such, any impact of RHOA on NS1 secretion was not considered further in this study. Compared to non-targeting siRNA controls, we identified 8 siRNA pools that inhibited NS1-NLuc secretion without significantly affecting cell viability. Figure 1C shows the extracellular NS1-NLuc -to-intracellular NS1-NLuc ratios (sNS1-NLuc / iNS1-NLuc), a measure of NS1 secretion efficiency, from the three independent siRNA screen repeats. Interestingly, the top three hits: COPA, COPB2, and COPG1, are three of the seven subunits of the coatomer protein complex I (COPI), a heptameric protein matrix that coats transport vesicles that operate within the early secretory pathway. COPI has recently been identified as being involved in various aspects of DENV biology (34, 35), however, this is the first study to directly implicate COPI as an important determinant of DENV NS1 secretion. Consistent with the results of our screen, previous studies have also identified several of our other top-ranking hits as host genes that are involved in orthoflavivirus NS1 biology. Intriguingly, dynamin-1 (DNM1) and early endosome antigen 1 (EEA1) have been shown to be involved in the internalisation of sNS1 (36), suggesting that these proteins may be involved in the bi-directional trafficking of secretion-destined NS1 and internalised sNS1. Similarly, valosin containing protein (VCP) has recently been shown to co-localise with NS1 protein in cells infected with Japanese encephalitis virus (37), raising the possibility that VCP-NS1 interactions may be a feature shared among orthoflaviviruses. The identification here of vav guanine nucleotide exchange factor 2 (VAV2) as a determinant of NS1 secretion may also relate to the reported involvement of VAV2 in DENV-induced inflammatory responses(38).

To validate the siRNA screen hits, a deconvolution screen was performed. Here, each of the four constituent siRNA duplexes from each pool was screened individually. While not identified as a hit in the screen, GBF1 was also included in this deconvolution screen due to its role as a master regulator of COPI vesicle formation (39). GBF1 is a guanine nucleotide exchange factor (GEF) that catalyses the GDP to GTP exchange on ADP-ribosylation factors (ARFs). GTP-activated ARFs then recruit preassembled heptameric COPI complexes to donor membranes to form COPI coated vesicles (29). For each of the nine gene targets, at least two siRNAs decreased NS1 secretion as inferred from the NS1-NLuc secretion ratio (Fig. 1D). Of the COPI components including GBF1, all but one of the individual siRNAs reduced the NS1-NLuc secretion ratio, providing further support that COPI machinery is involved in NS1 secretion. Taken together, several genes that encode components of the multi-subunit COPI complex and associated pathways were identified as critical factors involved in DENV NS1 secretion.

### NS1 secretion is reduced in COPI-silenced Huh-7.5 cells

Following identification of components of the COPI complex as putative determinants of efficient NS1 secretion, we next sought to confirm the effects of siRNA-mediated COPI gene knockdown and associated effects on NS1 secretion using wildtype infectious DENV2 and the related orthoflavivirus, Australian endemic West Nile virus Kunjin subtype (WNV/KUNV).

After validating the efficacy of our siRNAs in knockdown of the expression of their target mRNA in Huh-7.5 cells (Fig. 2A), we confirmed that COPI siRNA treatment successfully reduced target protein abundance by quantitative indirect immunofluorescence microscopy, using fluorescence intensity as a readout of protein abundance (Fig. 2B). Importantly, DENV-infected Huh-7.5 cell viability was largely unaffected by COPI siRNA treatment (Fig. 2C), as determined using an ATP-based cell viability assay. Infectious virus production was unaltered by COPI silencing (Fig. 2D), suggesting that COPI siRNA treatment does not impair DENV RNA replication, virion assembly, or virus egress when COPI gene knockdown is applied at 4 hours post-DENV infection. To assess the impact of COPI silencing on NS1 secretion, Huh-7.5 cells were infected with DENV for 4 hours and trypsinised and reverse transfected with siRNA pools targeting COPI components. At 24 hours post-infection (h.p.i.), cell culture media was replaced, and cell culture lysates and supernatants were harvested at 48 h.p.i. to assess intracellular and secreted NS1 abundance, respectively, by quantitative Western blot analysis (Fig 2E). Under each of the COPI siRNA treatments, DENV NS1 secretion was reduced as inferred from the NS1 secretion ratios (sNS1 / iNS1) relative to cells transfected with the non-targeting siRNA control (Fig. 2F), reflecting the results of the original siRNA screen. Similarly, a reduction in NS1 secretion was observed when COPI components were depleted in Huh-7.5 cells infected with WNV/KUNV (Fig 2G). Taken together, these data indicate that COPI components are required for efficient DENV NS1 secretion but are dispensable for infectious virus secretion. Further, the exploitation of COPI components to achieve NS1 secretion from human cells may be a conserved feature within the Orthoflavivirus genus.

**FIG 2.**
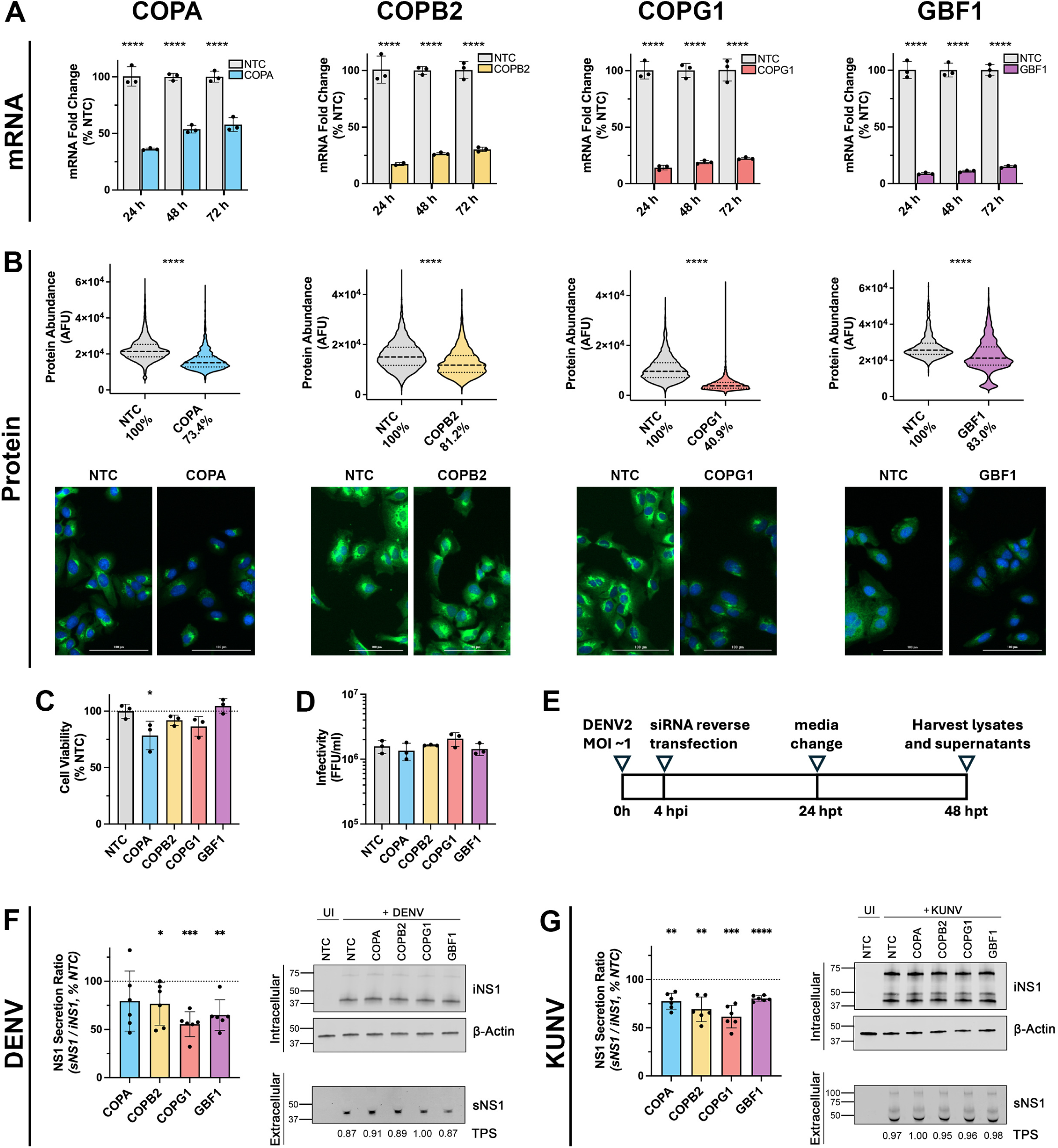
DENV and WNV/KUNV NS1 secretion is reduced in COPI silenced Huh-7.5 cells. (A) qRT-PCR analysis of COPI component mRNA levels in Huh-7.5 cells at indicated time points following siRNA treatment. Data are normalised to those of the RPLP0 housekeeping gene and expressed as a % of those of the non-targeting control (NTC) siRNA. Data are means + S.D., n = 3, one-way ANOVA. (B) Immunofluorescence microscopy-based quantitative analysis of COPI component protein abundance in Huh-7.5 cells following siRNA treatment. Huh-7.5 cells were reverse transfected with COPI siRNA pools or the NTC siRNA pool as indicated. At 48 h.p.t., cells were fixed and processed for indirect immunofluorescent labelling using anti-COPI antibodies (green) and nuclei were counterstained with DAPI (blue). Fluorescence intensity was measured for each cell to determine COPI protein abundance at the single cell level. Violin plots (with light smoothing) display median values (dashed lines) and quartile values (dotted lines) for each data set. Mean fluorescence intensity as a percentage of NTC are displayed on the x-axis. Cell numbers (n) COPA: NTC = 1902, COPA = 1630; COPB2: NTC = 6177, COPB2 = 3667; COPG1: NTC = 4146, COPG1 = 2153; GBF1: NTC = 1238, GBF1 = 1424. The statistical significance of differences between groups was determined using Welch’s t-test. (C) DENV-infected Huh-7.5 cell viability and infectious virus production is largely unaffected by COPI silencing. Huh-7.5 cells were infected with DENV2 (MOI ∼1) for 4 h and reverse transfected with siRNAs targeting the indicated COPI component or NTC. At 48 hours post-siRNA treatment, cell viability was measured using a CellTiter-Glo 2.0 cell viability assay (C) and virus-containing cell culture supernatants were recovered and processed to assess infectivity by focus forming assays (D). Data are means + S.D., n = 3 biological triplicates, one-way ANOVA. (E) Schematic overview of the experimental approach to assess the impact of COPI silencing on wildtype DENV2 or WNV/KUNV NS1 secretion. Huh-7.5 cells were infected with DENV2 or WNV/KUNV (MOI ∼1), trypsinised at 4 h.p.i., and reverse transfected with siRNAs targeting the indicated COPI components or NTC. At 24 hours post-siRNA treatment, cells were washed and media was replaced. At 48 hours post-siRNA treatment, cell culture supernatants and lysates were recovered to measure extracellular and intracellular NS1 levels, respectively, by quantitative Western blot analysis. (F – G) Quantification of NS1 abundance in cell culture supernatants and lysates by Western blot analysis, displayed as the secretion ratio of NS1 (sNS1 / iNS1) as a % of NTC. Data are means + S.D., n = 3 from two independent experiments, one-sample *t*-test. *p = <0.05, **p = <0.005, ***p = ≤0.0005, ****p = <0.0001.

### Heterologous expression of COPI variants suggests that certain variants and over-expression affect DENV NS1 secretion

Since siRNA-mediated depletion of COPI components reduced NS1 secretion in DENV-infected human cells, we next explored potential interactions between DENV2 NS1 and COPI components by immunofluorescence and confocal microscopy analysis. Huh-7.5 cells stably expressing GFP-tagged COPA, COPB2, or COPG1 cDNA were infected with DENV2 (MOI ∼0.1) for 24 hours prior to fixation, indirect immunofluorescent labelling using anti-NS1 and anti-GFP antibodies and confocal microscopy. While GFP-tagged COPI components displayed intense juxtanuclear and vesicular cytoplasmic staining patterns, consistent with Golgi-like localisation, these studies revealed infrequent colocalization between cytoplasmic NS1 foci and COPI vesicle marker staining (Figs. 3A, C and E), as highlighted by rare and incidental overlap of NS1- and COPI-associated peaks in line profile analyses (Figs. 3B, D and F). These results indicate that the vast majority of intracellular NS1 is spatially distinct from COPI components, suggesting that potential interactions between secretion-destined NS1 and COPI may be infrequent and/or transient in nature.

**FIG 3.**
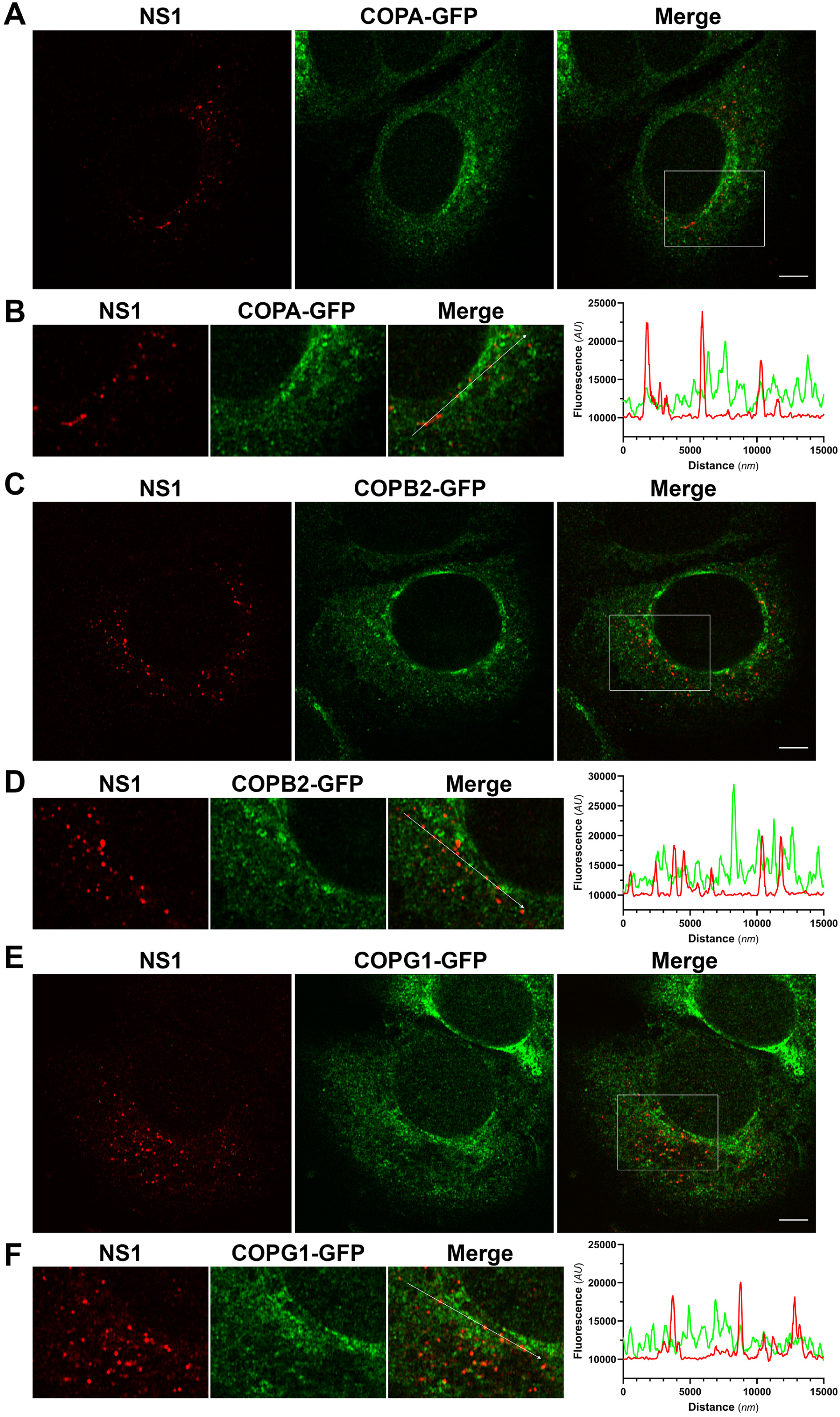
Confocal analysis of COPA, COPB2 and COPG1 localization reveals minimal and infrequent colocalization with NS1 protein in infected cells. Huh-7.5 cells stably expressing GFP-tagged COPA (A-B), COPB2 (C-D) or COPG1 (E-F) were infected with DENV2 (MOI ∼1) for 24 h prior to fixation and indirect immunofluorescent labelling using anti-NS1 (red) and anti-GFP (green) antibodies. Samples were processed for Airyscan confocal imaging and appropriate image processing, as described in Materials and Methods. Representative images, single central z-sections and merged images are shown, as indicated. Image panels in B, D and F depict ‘zoom insets’ taken from the indicated white box areas in Merge images in A, C and E, respectively. Graphs in B, D and F depict 15 µm line profiles for the lines indicated by white arrows in the corresponding ‘zoom inset’ Merge images. Scale bars, 10 µm.

Next, we sought to perturb the COPI pathway using an siRNA-independent approach. In line with previous studies (40, 41), our attempts to generate COPI component knockout cell lines using CRISPR/Cas9 failed to yield cells completely deficient in COPI component protein (data not shown), likely due to the established roles of these genes for optimal cell proliferation(42). However, medically relevant loss-of-function single nucleotide polymorphisms (SNPs) have recently been identified for COPA^-E241K^(43), COPB2^-R254C^(44), and COPG1^-K652E^(45). Accordingly, we investigated the impact of overexpression of wildtype and SNP variants of these genes on NS1 secretion efficiency. Importantly, COPA^-E241K^ is a dominant negative variant (43), therefore, heterologous cDNA expression should interfere with the proper functioning of the endogenously expressed wildtype protein. To date, no dominant negative SNP mutations have been identified for COPB2 or COPG1. Indeed, COPB2^-R254C^ and COPG1^-K652E^ are homozygous recessive mutations. Nevertheless, these variants are incorporated into COPI complexes resulting in impaired COPI function(45). To this end, GFP-tagged wildtype COPA, COPB2, and COPG1 cDNA constructs were generated and modified variants were generated by site-directed mutagenesis to incorporate these SNPs. To examine the effects of COPI^-WT^ or COPI^-SNP^ over-expression on NS1 secretion, independent of viral RNA replication and/or spread of infection, we employed the T7 RNA polymerase-driven pIRO-D expression system in which heterologously expressed T7 RNA polymerase drives expression of the DENV2 NS1-NS5 polyprotein and induces the formation of replication organelles that are morphologically indistinguishable to those of wildtype DENV infection(46). COPI^-WT^, COPI^-SNP^, or GFP-only control plasmids were co-transfected with pIRO-D into T7 RNA polymerase-expressing Huh-7.5 cells. At 18 hours post-transfection, cell culture lysates and supernatants were collected to assess the impact of COPI^-WT^ and COPI^-SNP^ cDNA over-expression on intracellular and secreted NS1 abundance by quantitative Western blot analysis. Despite substantial variability of iNS1 levels when either COPA^-WT^ or COPA^-E241K^ cDNA was expressed, sNS1 levels were relatively consistent within treatment groups (Fig. 4A). Interestingly, while COPA^-WT^ over-expression had no effect on sNS1 levels, expression of the COPA^-E241K^ construct increased sNS1 levels approximately two-fold. Modest increases in iNS1 abundance were observed when either COPB2^-WT^ or COPB2^-R254C^ constructs were over-expressed, however, an approximately two-fold increase in sNS1 abundance was observed in COPB2^-WT^ transfected cells (Fig. 4B). Similarly, modest increases in iNS1 were observed in cells transfected with either COPG1^-WT^ or COPG1^-K652E^ constructs, however, no effect was observed for levels of sNS1 (Fig. 4C). Collectively, the altered NS1 secretion profiles observed here for COPA^-E241K^ and COPB2^-WT^ suggests that allelic variants and/or altered expression levels of COPI components may enhance DENV NS1 secretion.

**FIG 4.**
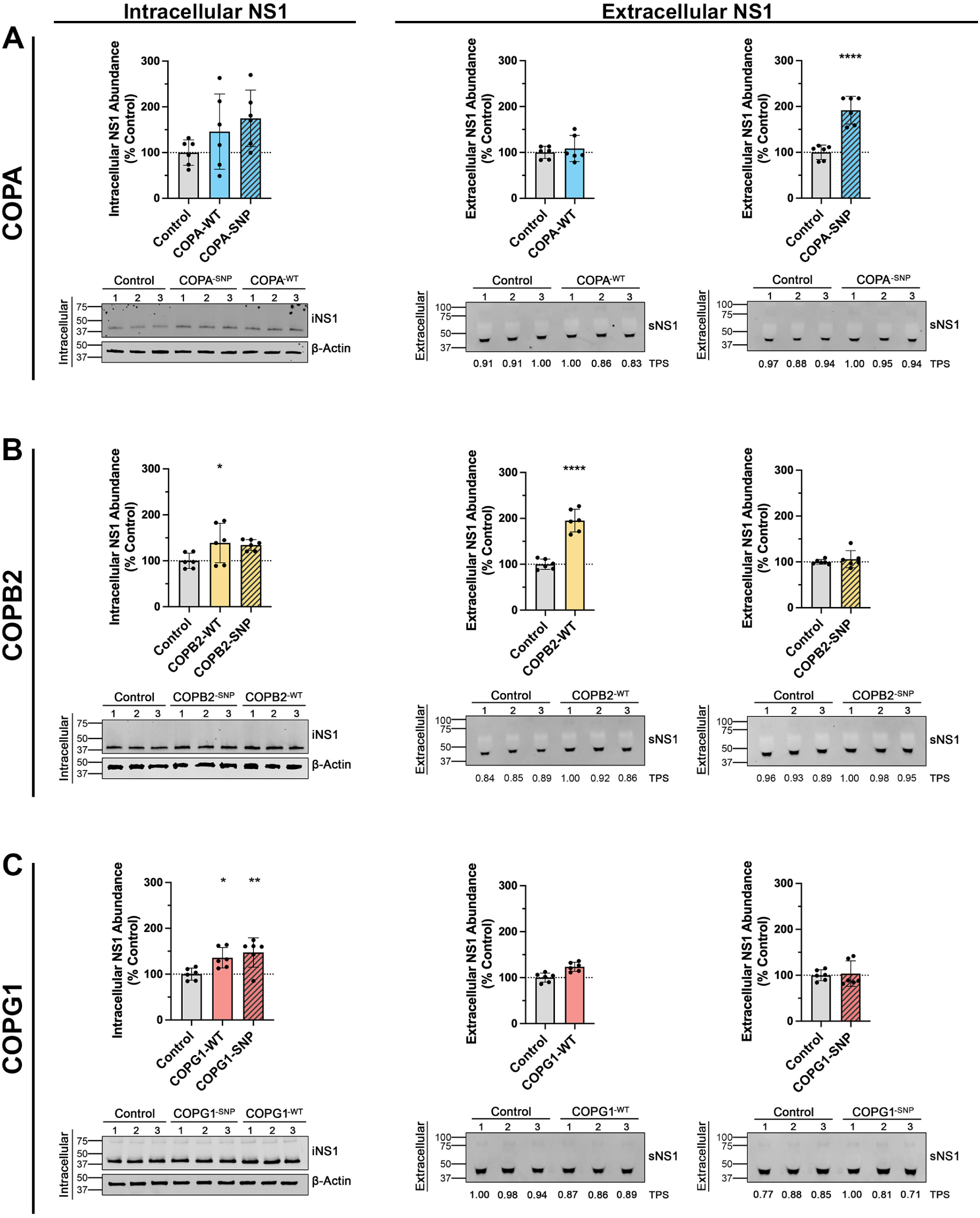
Heterologous expression of COPI variants indicates that certain variants and over-expression affect DENV2 NS1 secretion. Single nucleotide polymorphisms (SNPs) were introduced into GFP-tagged wildtype COPA, COPB2, and COPG1 cDNA expression constructs. T7 RNA polymerase-expressing Huh-7.5 cells were co-transfected with COPI expression constructs and a T7 RNA polymerase-driven DENV2 NS1-NS5 polyprotein expression system. At 18 hours post-transfection, cell culture supernatants and lysates were recovered to measure extracellular and intracellular NS1 levels, respectively, by quantitative Western blot analysis. Data are means + SD, n = 3 from two independent experiments, one-way ANOVA, *p = <0.05, **p = <0.01, ***p = <0.005, ****p = <0.0001.

### NS1 secretion is reduced in Golgicide A-treated Huh-7.5 cells

Golgicide A (GCA) is a potent and specific inhibitor of GBF1 catalytic activity that acts by binding to the GBF1-Arf-GDP protein-protein interface, preventing the Arf-GDP/GTP exchange (47). This results in COPI dissociation from Golgi membranes, prevention of COPI vesicle formation, disassembly of the Golgi, and swelling of the ER (47). GBF1 has been demonstrated to play a variety of roles in many RNA virus lifecycles (48), and much of this information has been elucidated by the use of GCA or the related multi-ArfGEF inhibitor brefeldin A (BFA). Importantly, time of addition studies have shown that these compounds influence multiple aspects of the orthoflavivirus life cycle. When applied to WNV/KUNV infected cells during the 12-16 h latent phase of infection (49), BFA inhibits the formation of virus induced membrane structures (50), and severely impairs viral protein production and infectious virus release (51). However, when added late in infection, the virus induced membrane structures are relatively stable (50), and only minor effects on protein synthesis were observed (51). Comparatively, GCA pulse-chase studies in DENV-infected cells indicate that despite having no impact on DENV internalisation, intracellular viral RNA abundance is significantly reduced when GCA is applied in the first 12 h of infection, reduced to a lesser extent when applied at 12 h.p.i., but unaffected when applied at 24 h.p.i. (52). Indeed, the extracellular accumulation of orthoflavivirus NS1 has been shown to be decreased in orthoflavivirus-infected mammalian cells when treated with high concentrations BFA or GCA at 1 h.p.i. (53). However, given the inhibitory effects of these compounds on viral RNA replication, protein synthesis, and infectious virus production when administered early during infection, this is not surprising. As such, to mitigate the inhibitory effect of GCA on DENV RNA replication, we examined the impact of GCA treatment on NS1 secretion in DENV-infected cells when GCA is administered late in infection.

To investigate the effect of GCA on various aspects of DENV biology, Huh-7.5 cells were infected with DENV for 24 hours. At 24 h.p.i., cell culture media was replaced with GCA-supplemented media (0 – 5 µM) and returned to culture for a further 18 hours prior to analysis. First, we tested the effect of GCA treatment on DENV-infected Huh-7.5 cell viability using an ATP-based cell viability assay. No significant effects on cell viability were observed at GCA concentrations ≤ 5 µM (Fig. 5A). Increasing concentrations of GCA from 1 to 5 µM did, however, reveal a dose-dependent reduction of infectious virus production (Fig. 5B), accompanied by increases in intracellular viral RNA abundance (Fig. 5C). The use of a *Renilla* luciferase-encoding subgenomic replicon confirmed that this increase in intracellular viral RNA abundance was not the result of changes to viral RNA replication (Fig. 5D). These results suggest that GCA-mediated GBF1 inhibition does not influence DENV genome replication but, instead, impedes infectious DENV particle release when GCA is applied to cells after 24 hours of DENV infection. To assess the impact of GCA on NS1 secretion, cell culture lysates and supernatants were recovered from DENV-infected, GCA-treated Huh-7.5 cells. A decrease in NS1 secretion, as inferred from the NS1 secretion ratio (sNS1 / iNS1), was observed in cells treated with 5 µM GCA, indicating that GCA reduces NS1 secretion from DENV-infected Huh-7.5 cells (Fig. 5E). Similar reductions in NS1 secretion were observed for WNV/KUNV-infected Huh-7.5 cells treated with 5 µM GCA (Fig. 5F). This 5 µM GCA-induced reduction in NS1 secretion appears to be independent of changes to Golgi morphology, as no apparent differences in NS1 and the Golgi marker GM130 staining patterns were observed by confocal immunofluorescence microscopy, and co-localisation analysis indicated no significant impact on NS1 co-localisation with GM130 (Fig. 5G). Collectively, these findings indicate that, when GCA is applied at 24 hours post-infection, the catalytic activity of GBF1 is dispensable for DENV genome replication but is required for efficient virus secretion and NS1 secretion.

**FIG 5.**
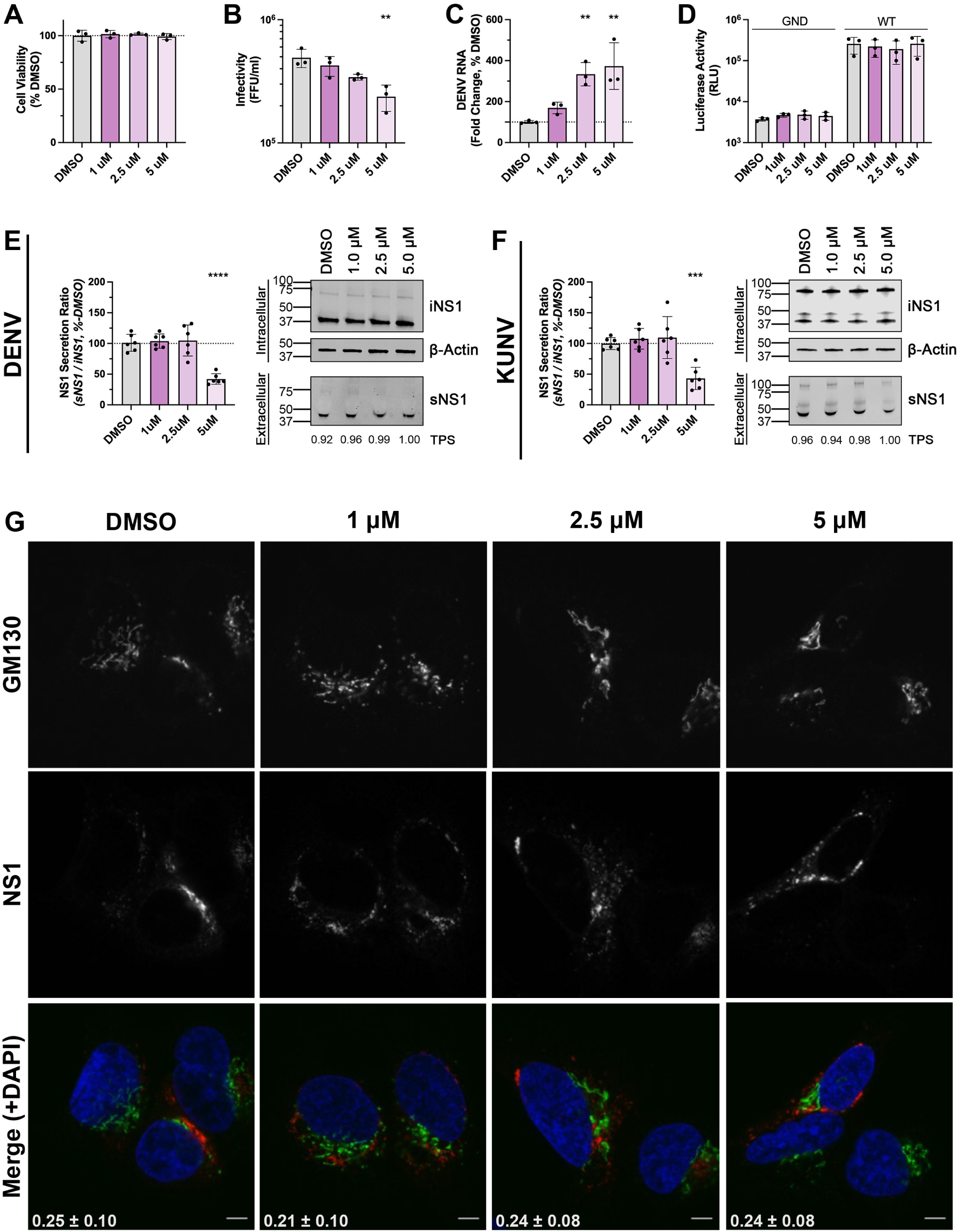
DENV2 and WNV/KUNV NS1 secretion is reduced in Golgicide A-treated Huh-7.5 cells. (A-C) Impact of Golgicide A (GCA) on DENV biology. Huh-7.5 cells were infected with DENV2 (MOI ∼1). At 24 hours post-infection, cells were washed and cultured for a further 18 h in media supplemented with increasing concentrations of GCA or DMSO control. At 18 hours post-GCA treatment, cell viability was measured using CellTiter-Glo 2.0 viability assay (A), virus-containing cell culture supernatants were recovered and processed to assess infectivity by focus forming assay (B), and total cellular RNA was collected for qRT-PCR analysis of DENV2 viral RNA levels. For qRT-PCR analysis, data are normalised to the RPLP0 housekeeping gene and expressed as a % of non-targeting control (NTC) siRNA-treated mean values (C). Data are means + SD, n = 3 biological triplicates. (D) Golgicide A does not impact DENV RNA replication. Huh-7.5 cells were transfected with *in vitro* transcribed (IVT) RNA for a DENV2 subgenomic reporter replicon sg-DVs-R2A (WT, or replication-deficient GND control). At 4 hours post-transfection, cells were lysed (4 h timepoint) or cultured in GCA at the indicated concentration or DMSO carrier control. At 18 h post-GCA treatment, cell lysates were prepared and luciferase activities were determined as a surrogate marker for viral RNA replication. Data are means + SD, n = 3 biological triplicates. (E – F) Quantification of NS1 abundance in cell culture supernatants and lysates, displayed as secretion ratio of NS1 (sNS1 / iNS1) as a % of DMSO. Data are means + SD, n = 3 from two independent experiments, one-way ANOVA, **p= <0.005, ***p= <0.001, ****p = <0.0001. (F) Localisation of NS1 with respect to the Golgi marker GM130. Huh-7.5 cells were cultured as shown in E. At 18 hours post-GCA treatment, cells were fixed and stained for indirect immunofluorescent labelling using mouse anti-NS1 and rabbit anti-GM130 primary antibodies, followed by AlexaFluor 555-conjugated anti-mouse IgG (red) and AlexaFluor 488-conjugated anti-rabbit IgG (green). Samples were counterstained with DAPI and analysed by confocal fluorescence microscopy. Yellow in the merged images indicates co-localisation. Pearson’s co-localisation coefficients are shown in white in the merged images (means + SD, n = >30 cells). Scale bars are 10 µM.

## DISCUSSION

To identify host proteins that are involved in the secretion of DENV NS1, we performed a customised membrane-trafficking siRNA screen in human Huh-7.5 hepatoma cells infected with the DENV2-NS1-NLuc reporter virus. Our screen revealed the coatomer protein complex I (COPI) subunits COPA, COPB2, and COPG1, as the top three hits whose depletion reduced extracellular levels of NS1, thus heavily implicating COPI as an important determinant of DENV NS1 secretion.

COPI is a highly conserved protein complex that coats transport vesicles that shuttle protein and lipid cargo between cellular compartments. The complex consists of seven coatomer subunits (*⍺*, *β*, *β*^r^, *δ*, *ɛ*, *γ*, *ζ*) (54). Mechanistically, COPI vesicle formation requires GBF1-catalysed hydrolysis of GDP for GTP on ADP-ribosylation factors (Arf) (39, 55). Activated Arfs then recruit preassembled cytosolic heptameric COPI complexes to a donor membrane(56). The continued recruitment of COPI complexes to the nascent vesicle results in membrane destabilisation and ultimately culminates in vesicle scission (29). The newly formed COPI coated vesicle, complete with membrane bound and luminal cargo is then disseminated to its target acceptor membrane location (57). The best categorised role of COPI coated vesicles is their involvement in the bi-directional trafficking of protein and lipids within the early secretory pathway (30). COPI coated vesicles function in intra-Golgi trafficking mediating anterograde and retrograde transport (58, 59). They also mediate Golgi-to-ER recycling of escaped ER-resident proteins, thus maintaining the structural and functional integrity of these organelles(60). Several studies have implicated COPI components as performing a role in endosomal transport and function (61–64). More recently, COPI has been demonstrated to perform roles in a wealth of processes including lipid metabolism (65), autophagy (66), mRNA localisation (67), nuclear envelope disassembly (68), and neurogenesis (69, 70). Regarding COPI vesicle regulators, GBF1 is well documented as being involved in multiple aspects of orthoflavivirus replication (reviewed in (48)). In addition, several Arfs have also been shown to play overlapping and redundant roles in DENV biology (35, 71). While these components regulate COPI vesicle formation, it must be noted that they have multiple effectors (72–74). However, given that the effects of GBF1 and Arf inhibition on orthoflavivirus biology can be phenocopied by COPI component depletion (35), this strongly suggests that COPI is involved in multiple aspects of the orthoflavivirus life cycle. Crucially, a recent study by Iglesias et al. (35) demonstrated that DENV utilises COPI for the trafficking of capsid protein between the ER and lipid droplets, highlighting that the exploitation of COPI machinery by DENV is not limited to the canonical role of COPI in the secretory pathway.

Given the diverse roles of COPI and its regulators in orthoflavivirus biology, to focus specifically on NS1 secretion while minimising pleiotropic effects, we concentrated our attention towards perturbing the COPI pathway at later stages of infection. Using this approach, we confirmed that siRNA-mediated depletion of COPI did not significantly impact intracellular DENV viral RNA replication or infectious virus production. Importantly, however, COPI depletion did result in a decrease in extracellular levels of NS1, coincident with an increase in the intracellular levels of NS1 for both DENV and WNV/KUNV, indicating that COPI siRNA mediated depletion impairs the efficient secretion of NS1 for multiple orthoflaviviruses. While the modest levels of NS1 secretion inhibition observed here may reflect incomplete COPI protein knockdown, as preassembled heptameric COPI complexes are relatively stable and display a half-life of ∼28 h in mammalian cells(75), these results also suggest the possible existence of multiple mechanisms that may be exploited by orthoflaviviruses to achieve NS1 secretion from human cells.

Intriguingly, pathogenic COPI SNP variants have been implicated in causing disease phenotypes that reflect those associated with orthoflavivirus complications, including arthritis(43), haemorrhage(43), microcephaly(44, 76) and dysregulation of immune system(45)). By overexpressing wildtype or deleterious COPI alleles in cells co-transfected with a replication-independent DENV non-structural protein expression vector, we were able to assess the effect of COPI perturbation on NS1 secretion independently from genome replication and infectious virus spread. The increase in the secretion efficiency of NS1 from cells transfected with the wildtype COPB2 construct suggests that the availability of COPB2, but not COPA nor COPG1, protein may represent a bottleneck and be a limiting factor in NS1 secretion. Unexpectedly, the overexpression of the dominant-negative mutant COPA^-E241K^ appeared to enhance NS1 secretion. This COPA allele contains a mutation in the WD40 domain which causes defects in Golgi-to-ER trafficking and leads to increases in ER stress(43). Whether the observed increase in NS1 secretion is a direct effect of COPA^-E241K^ expression remains an open question. It is possible that NS1 secretion is achieved via a non-canonical COPI function whereby the WD40 domain of COPA may be dispensable or even inhibitory to NS1 secretion. Alternatively, NS1 secretion may be favoured under conditions of enhanced ER stress brought about by COPA^-E241K^ overexpression. Nonetheless, how the overexpression of the COPA^-E241K^ variant enhances NS1 secretion warrants further investigation. Similar to the effects of COPA^-E241K^, COPB2^-R254C^ and COPG1^-K652E^ also show defects in Golgi-to-ER trafficking and aberrant cellular responses (44, 45, 77). They are, however, not dominant-negative and as such their heterologous expression in a wildtype COPI background may have masked any potential effect on NS1 secretion. Despite our apparent inability to generate COPI component knockout cell lines by CRISPR/Cas9 technology, the use of genome editing to introduce these genetic variants in the place of wild-type genes offers an attractive approach to further explore the emerging roles of COPI in orthoflavivirus biology.

To functionally inhibit the formation of COPI vesicles, we employed the small molecule inhibitor Golgicide A (GCA). While it is well documented that GCA mediates a variety of effects on orthoflavivirus biology(48), most studies have utilised this compound at early timepoints during infection. Here, we functionally inhibited GBF1 using GCA at a later stage of infection and found that GCA reduces infectious DENV production in a dose-dependent manner. Consistent with a GCA-mediated defect in infectious virus production, we observed a concomitant increase in the intracellular levels of DENV viral RNA. The use of a DENV subgenomic replicon verified that GCA was not impacting viral RNA genome replication, confirming that GCA acts to prevent the release of infectious DENV virions. Interestingly, while virion secretion was reduced by GCA treatment in a dose-dependent manner, NS1 secretion was only observed to be reduced at the highest dose applied. Importantly, our results indicate that infectious DENV production is more sensitive than NS1 secretion to the effects of GCA-mediated GBF1 inhibition. These results provide further support to the conclusion that multiple mechanisms, both GBF1-dependent and GBF1-independent, may be exploited by orthoflaviviruses to achieve NS1 secretion from human cells.

Many genetic, biochemical, and imaging studies have been performed to interrogate NS1 secretion biology and these have been integral in defining sNS1 structure and key functional residues. It is widely assumed that DENV NS1 is secreted from mammalian cells via the canonical secretion pathway (78). Multiple studies have indicated that DENV NS1 is translated into the ER as a soluble monomer and decorated with high-mannose moieties at N130 and N207 (6, 7, 79, 80). NS1 monomers rapidly homodimerize to form a partially hydrophobic NS1 dimer, the predominant intracellular form (6, 7). Membrane-associated NS1 dimers are proposed to preferentially localise to the sites of nascent lipid droplets on the luminal side of the ER (12), or to cholesterol-rich microdomains within the Golgi apparatus (3, 4, 11, 12). This has been suggested as a mechanism to concentrate NS1 dimers, with three dimers thought to come together to pinch off from the membrane, converting them into a soluble hexamer and collecting the lipid component that fills the hexamer’s central cavity(12). While not a strict prerequisite for NS1 secretion(80–82), the secreted form of NS1 exhibits a complex-type glycan at N130(80). It is proposed that this additional processing of the N130 glycan occurs following ER-to-Golgi translocation given that, in uninfected cells, the machinery responsible for this maturation resides in the Golgi(6). Secretion-destined NS1 is understood to then traffic from the trans-Golgi network to the plasma membrane where it exits the cell as a hexameric glycolipoprotein (78, 83). This study confirms that COPI components are important determinants of NS1 secretion and this is compatible with the canonical secretion pathway model. However, the results of our GCA experiments are particularly intriguing. It is well established that DENV virions are matured as they traffic through the secretory pathway prior to being released from the cell as fully infectious virions (84). While the relatively low concentrations of GCA employed here revealed a dose-dependent reduction in infectious DENV production, a reduction in NS1 secretion was only observed at the highest dose (5µM). Moreover, despite a dramatic decrease in NS1 secretion in cells treated with 5µM GCA, our confocal microscopy studies showed no significant impact of GCA on NS1 and Golgi marker GM130 co-localisation. Given the additional and emerging roles of COPI beyond intra-Golgi and Golgi-to-ER trafficking, and the demonstration that DENV exploits a non-canonical role of COPI to traffic capsid protein, alternative roles of COPI involvement in NS1 secretion warrant consideration. Potential sites of COPI involvement in NS1 secretion are shown in Figure 6. Future studies further interrogating the contribution that COPI coated vesicles, their activators, and their vesicle constituents play in orthoflavivirus NS1 secretion will be integral to defining the role(s) of COPI in NS1 secretion and may provide additional targets for NS1-specific anti-orthoflaviviral therapies.

**FIG 6.**
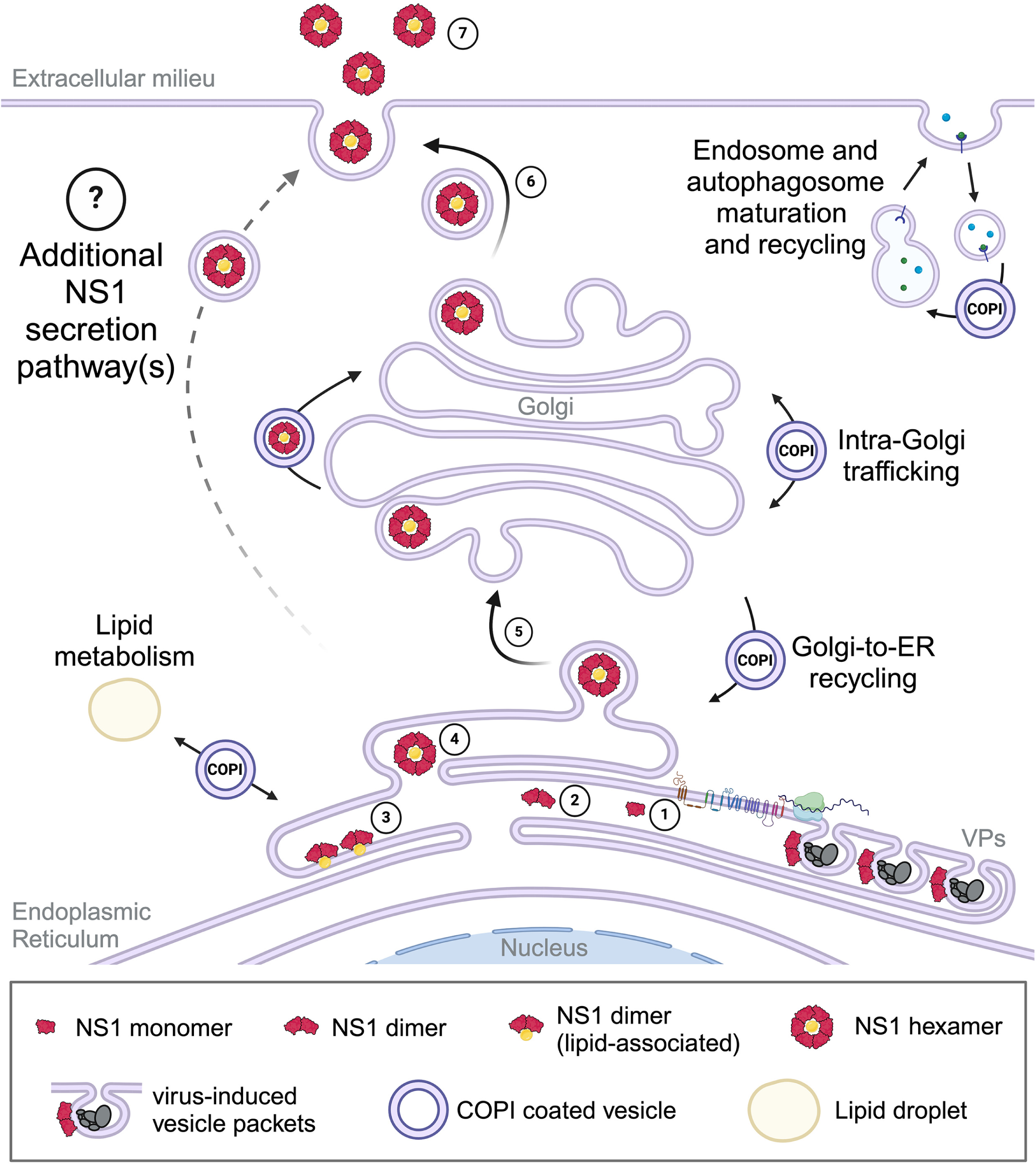
Potential sites of COPI involvement in DENV NS1 secretion. (1) DENV NS1 is translated into the ER as a soluble monomer and modified by the addition of high-mannose glycans at N130 and N207. (2) Soluble monomers rapidly homodimerize to form a partially hydrophobic and membrane-associated dimer, the predominant intracellular NS1 form that plays a critical role in viral genome replication (vesicle packet (VPs). (3) Membrane-associated NS1 dimers are proposed to concentrate at sites of nascent lipid droplets within the ER or cholesterol-rich microdomains within the Golgi. (4) Three membrane-associated dimers come together and pinch off from the membrane to form a soluble NS1 hexamer that is stabilised by a central lipid component. (5) Secretion-destined NS1 is proposed to traffic from the ER to the Golgi for additional processing of the N130 moiety to a complex-type glycan. (6) NS1 is proposed to be dispatched from the trans-Golgi network to the plasma membrane where it is released into the extracellular environment. (7) Secreted NS1 promotes viral propagation and contributes to dengue disease pathogenesis through a variety of pathways. Potential sites of COPI participation in NS1 secretion are shown (see Discussion). Created with BioRender.

## MATERIALS AND METHODS

### Cell culture

Huh-7.5 cells (85) were generously provided by Charles M. Rice (Rockefeller University, New York, USA). Huh-7.5+Fluc cells stably expressing firefly luciferase have been previously described (86). Huh-7.5+T7 RNApol cells stably expressing T7 RNA polymerase have been previously described (87). All cells were maintained as previously described(87).

### Antibodies and chemicals

Mouse anti-NS1 monoclonal antibody (MAb) 4G4 was generously provided by Jody Peters and Roy Hall (University of Queensland, Brisbane, Australia) (88) or purchased from Mozzy Mabs (University of Queensland, Brisbane, Australia). Mouse anti-capsid MAb 6F3.1 was kindly provided by John Aaskov (Queensland University of Technology, Brisbane, Australia) (89). The mouse anti-Envelope monoclonal antibody was prepared from the hybridoma cell line D1-4G2-4-15 (4G2) purchased from ATCC and maintained as previously described (90). Mouse anti-β-actin MAb (AC-15) was purchased from Sigma-Aldrich. Mouse anti-COPA MAb (sc-398099) was purchased from Santa Cruz Biotechnology. Rabbit anti-COPB2 polyclonal antibody (PAb) (ab2899) and rabbit anti-GBF1 PAb (ab86071) were purchased from Abcam. Rabbit anti-COPG1 PAb (PA5-65194) and rabbit anti-GFP PAb (A-11122) were purchased from Thermo Fisher Scientific. Rabbit anti-GM130 MAb (D6B1) was purchased from Cell Signaling Technology. Alexa Fluor 488- and 555-conjugated anti-mouse IgG and anti-rabbit IgG secondary antibodies were purchased from Thermo Fisher Scientific. Fluorescent stain 4’,6-diamidino-2-phenylindole (DAPI) was purchased from Sigma-Aldrich. Revert™ 700 Total Protein Stain and IRDye® 800 CW goat anti-mouse secondary antibody for Western Immunoblotting were purchased from LI-COR Biosciences. Golgicide A was purchased from Sigma-Aldrich and dissolved in dimethyl sulfoxide DMSO to 10 mM, aliquoted and stored at -20°C.

### Virus plasmids and virus propagation

Plasmid pFK-DVs containing a full-length DENV2 genome (strain 16681), a pFK-DVs subgenomic replicon derivative encoding a *Renilla* luciferase construct (pFK-sgDVs-R2A), a pFK-sgDVs-R2A replication-deficient NS5 mutant derivative (pFK-sgDVs-GND-R2A), and a replication-independent T7-driven DENV2 non-structural protein 1-5 expression system (pIRO-D) were generously provided by Ralf Bartenschlager (University of Heidelberg, Heidelberg, Germany) (46, 91). pFK-DVs-NS1-NLuc bearing a NanoLuc luciferase tag within NS1 was created as previously described (9). Infectious DENV2 stocks were generated following *in vitro* transcription of DENV2 RNA using mMessage mMachine SP6 RNA polymerase (Thermo Fisher Scientific) using *Xba*I-linearised plasmid as template DNA and transfection of Huh-7.5 cells with purified RNA using DMRIE-C (Thermo Fisher Scientific) transfection reagent, as previously described (90). WNV/KUNV was generously provided by Karla J. Helbig (La Trobe University, Melbourne, Australia).

### Generating GFP-tagged wildtype and SNP variant COPI cDNA expression constructs and associated stable cell lines

To generate GFP-tagged wildtype COPB2 and COPG1 cDNA expression constructs, FLAG-tagged COPB2 and COPG1 cDNA ORF clone plasmids (GenScript: Cat No. Ohu04585 and Cat No. Ohu29546D, respectively) served as cDNA template. For COPA cDNA, total cellular RNA was isolated from Huh-7.5 cells using NucleoZOL (Macherey-Nagel), as per manufacturer’s instructions. First-strand cDNA was synthesised using M-MLV Reverse Transcriptase (Promega) in conjunction with random hexamer primers as per manufacturer’s instructions. COPA, COPB2, and COPG1 cDNA was PCR amplified using gene specific primers (Supplementary Table 2) and Q5® High-Fidelity DNA Polymerase (New England Biolabs). Gel purified (Macherey-Nagel) amplicons were assembled as in-frame fusions with emGFP PCR product into a pLenti6/V5-D-TOPO vector using NEBuilder HiFi DNA Assembly (New England Biolabs). cDNA sequences were confirmed by Sanger Sequencing (Australian Genome Research Facility, Adelaide, Australia).

For SNP incorporation, GFP-tagged COPI cDNA constructs were modified by QuikChange II site-directed mutagenesis (Agilent) in conjunction with SNP-specific primers (Supplementary Table 2) and confirmed by Sanger sequencing. Exact cloning details for these constructs are available on request.

To generate stable Huh-7.5 cell lines expressing emGFP-tagged COPA, COPB2 and COPG1, the respective pLenti6/V5-D-TOPO plasmids containing these cassettes were packaged into lentiviral vectors and used to transduce Huh-7.5 target cells as described previously (Eyre NS, Drummer HE, Beard MR PLoS Pathogens 2010). Following selection with 5 µg/ml of Blasticidin HCl (Thermo Fisher Scientific) for approximately 2 weeks, emGFP-positive cells were sorted using a BD FACSAria III Fusion Flow Cytometer (BD Biosciences) at the Flinders University Flow Cytometry Facility and expanded and maintained as polyclonal cell lines.

### siRNA library screening

The siRNA library comprised a commercially available library targeting membrane trafficking proteins (Human ON-TARGETplus siRNA Library-Membrane trafficking – SMARTpool, Dharmacon cat# G-105500) and 37 additional siRNA SMARTpools (Dharmacon, Horizon Discovery) targeting previously identified proviral host factors that may be manipulated by NS1 (32, 92). Each siRNA SMARTpool consists of four siRNAs targeting the same gene. A scrambled non-targeting control (NTC) siRNA served as a negative control. siRNAs targeting Firefly luciferase (FLuc) and NanoLuc luciferase (NLuc) served as controls for cell viability and inhibition of DENV replication, respectively. The siRNA library was prepared at 1 µM in siRNA buffer (Dharmacon cat# B-002000-UB-100), pre-arrayed in 96-well plates at a volume of 4 uL/well, as appropriate for individual screening experiments and stored at -80°C until further use. siRNA screening and screen data analysis was performed at Cell Screen SA (Flinders Centre for Innovation in Cancer, Flinders University, Australia).

For a schematic representation of the experimental setup, see Fig. 1B. Huh-7.5+FLuc cells were seeded into 75cm^2^ flasks at 1.56 × 10^6^ cells/flask. The following day, cells were transfected with DENV2-NS1-NLuc *in vitro* transcribed (IVT) RNA using DMRIE-C for 3 h before transfection reagent was replaced with complete media. At 48 hours post-transfection (h.p.t.), DENV2-NS1-NLuc RNA-transfected cells were trypsinised and reverse transfected with the siRNA SMARTpool library at a final concentration of 40 nM in 96-well plates. For this, prearrayed 96-well plates containing 4uL of 1uM siRNA SMARTPool siRNAs were incubated with 15.7uL OptiMEM and 0.3uL DharmaFECT4 (Horizon Discovery) for 20 mins before DENV2-NS1-NLuc transfected cells were added at 1.25 × 10^4^ cells/well/80uL. After 3 hours of incubation, siRNA transfection reagent-containing media was replaced with complete media (100 uL/well). For each experiment, each siRNA SMARTPool was transfected in triplicate and 3 independent experimental replicates were performed. At 48 hours post-siRNA transfection, cell culture supernatants were collected, clarified by centrifugation (500 x g, 5 min, at 15°C) and mixed 1:1 with 2× passive lysis buffer (Promega cat #E1941). Cell monolayers were washed in PBS and lysed in 1× passive lysis buffer. Cell lysates and lysed-supernatants were stored at -20°C until further use. To assess the impact of siRNA SMARTpool treatment on intracellular and extracellular NS1 abundance, samples were assayed using the Nano-Glo Dual-luciferase reporter (NanoDLR) assay (Promega), according to manufacturer’s recommendations. Cell lysate or lysed-supernatant was mixed with OneGlo reagent (Promega) and incubated for 45 min at RT prior to FLuc (cell viability) luminescence quantification using a PerkinElmer Ensight plate reader. Following FLuc luminescence measurements, NanoDLR Stop&Go reagent was added, mixed, and incubated for 45 min at room temperature (RT) prior to NS1-associated NLuc luminescence quantification. NLuc-associated luminescence values and FLuc-associated luminescence values for each siRNA treatment were expressed as a percentage of corresponding average NTC siRNA-associated values. Means, S.D., and % CV were calculated on these normalised values and the test siRNA were scored to identify hits as detailed in the Supplemental Material.

A deconvolution screen was performed on the 8 criteria-matching hits and GBF1. Here, each of the four individual siRNA duplexes comprising each siRNA SMARTPool was assayed in triplicate. In this screen cell viability was additionally measured using CellTiter-Blue (Promega), as per manufacturer’s instructions, using a PerkinElmer Ensight plate reader. Transfections and luciferase assays were performed as described above. Data analysis and hit scoring was performed as described above and in the Supplemental Material.

### siRNA treatment of flavivirus infected cells

Huh-7.5 cells were grown in 75 cm^2^ flasks (5 × 10^6^ cells/flask) and cultured overnight. The following day, cells were infected with DENV2 or KUNV (MOI ∼1) or mock infected. At 4 hours post-infection (h.p.i.), cells were trypsinised and reverse transfected with COPI siRNA SMARTpools at a final concentration of 40 nM in 12-well plates. For this, a transfection mix comprised of 2 µL of 20 µM siRNA SMARTPool, 195 µL OptiMEM and 3 µL Dharmafect4 was incubated at RT for 20 mins before DENV- or KUNV-infected cells were added at 1.8 × 10^5^ cells/well/800µL. After 3 hours of incubation, siRNA transfection reagent-containing media was replaced with complete media (1mL/well). At 24 h.p.i., cells were washed and media was replaced with complete DMEM (400 µL/well). At 48 h.p.i., cell culture lysates and supernatants were recovered to measure intracellular and extracellular NS1 abundance, respectively. In parallel DENV-infected plates, total cellular RNA and cell culture supernatants were collected to measure viral RNA levels, host RNA expression and infectious virus production, respectively.

### COPI cDNA and pIRO-D expression vector co-transfection

Huh-7.5+T7 RNApol cells were seeded in 12-well plates at 1.2 × 10^5^ cells/well and cultured overnight in complete DMEM without puromycin. The following day, cells were co-transfected with 0.5 µg of each plasmid pIRO-D and COPI cDNA well using Lipofectamine 3000 (Thermo Fisher Scientific) according to manufacturer’s instructions and transfection reagent-containing media was replaced with complete DMEM at 3 h.p.t.. At 18 h.p.t., cell culture lysates and supernatants were collected for Western blot analysis.

### Golgicide A (GCA) treatment of flavivirus infected cells

Huh-7.5 cells were seeded into 75 cm^2^ flasks at 5 × 10^6^ cells/flask and cultured overnight. The following day, cells were infected with DENV2 or KUNV (MOI: ∼1) or mock infected. At 4 h.p.i., cells were washed, trypsinised and re-seeded into 12-well plates at 1 × 10^5^ cells/well for Western blotting, qRT-PCR or focus forming assays (FFA) or 96-well plates at 1 × 10^4^ cells/well for cell viability assays and returned to culture in complete DMEM. At 24 h.p.i., cells were washed, and cultured in complete DMEM supplemented with the indicated concentrations of GCA or DMSO carrier control, as indicated. At 18 hours post-GCA treatment, samples were harvested for Western blot, RT-qPCR, FFA, and cell viability analysis, as indicated.

### Quantification of mRNA and viral RNA by RT-qPCR

Total cellular RNA was extracted from near-confluent cells in 12-well plates using NucleoZOL (Macherey-Nagel), according to manufacturer’s instructions. Both first-strand complementary DNA (cDNA) synthesis and RT-qPCR was performed using the Luna Universal One-Step RT-qPCR Kit (New England Biolabs) in 384-well plates using a QuantStudio 7 Flex (Life Technologies) or CFX Opus (Bio-Rad) thermal cycler according to the manufacturer’s recommendations. For each sample and each primer pair, 10 µL reactions were prepared in technical duplicate, each containing 5 ng of total RNA, 0.2 µL of each primer at 20 µM (0.4 µM final concentration), 0.5 µL Luna WarmStart RT Enzyme Mix and DNase/RNase-Free water. Melt curve analysis was performed using the qPCR instrument default settings. mRNA or viral RNA levels were expressed as a percentage of the experimental control (NTC or DMSO, as appropriate) following normalisation to RPLPO mRNA, using the threshold cycle (ΔΔCT) method. Primer sequences are available upon request.

### Quantification of subgenomic DENV RNA replication by luciferase assay

Subgenomic RLuc-encoding DENV2 replicons were subjected to *in vitro* RNA transcription as previously described(90). Huh-7.5 cells were seeded in 12-well plates at 1 × 10^5^ cells/well and cultured overnight. The following day, cells were transfected with IVT RNA using DMRIE-C (Thermo Fisher Scientific) according to the manufacturer’s instructions. At 3 h.p.t., transfection reagent-containing media was replaced, and cells were returned to culture in complete DMEM. At 24 h.p.t., cells were washed and returned to culture in complete DMEM supplemented with increasing concentrations of GCA or 0.1% (v/v) DMSO carrier control alone, as indicated. At 18 hours post-GCA treatment, cell culture monolayers were lysed in Renilla Luciferase Lysis Buffer (Promega). Samples were mixed with Renilla Luciferase Assay Reagent (Promega) according to the manufacturer’s instructions, and luminescence was determined using a Cytation 5 Multimode Reader equipped with a Dual-Reagent Injector Module (BioTek).

### Quantification of protein knockdown by indirect immunofluorescence microscopy

Huh-7.5 cells were reverse transfected in 12-well plates at 1 × 10^5^ cells/well with COPI or NTC siRNA SMARTpools at a final concentration of 40 nM. At 3 h.p.t., transfection media was replaced with complete DMEM and cells were returned to culture. At 24 h.p.t., cells were trypsinised and re-seeded into a 96-well black-walled imaging plates (PerkinElmer PhenoPlate-96) at 1 × 10^4^ cells/well and returned to culture. At 48 h.p.t. cells were washed with PBS prior to fixation using ice-cold acetone:methanol (1:1), for 5 min at 4°C and then blocked using 5%(w/v) BSA in PBS for 30 min at RT. Samples were labelled with indicated primary antibodies anti-COPA (1:50), anti-COPB2 (1:100), anti-COPG1 (1:100), or anti-GBF1 (1:100) diluted in 1% BSA in PBS at 4°C overnight. Samples were then washed three times in PBS before incubation for 2 h in the dark at 4°C with Alexa Fluor 488-conjugated anti-rabbit IgG or Alexa Fluor 488-conjugated anti-mouse IgG antibodies (Thermo Fisher Scientific), as appropriate, diluted to 1:500 and 1:2000, respectively. Samples were then washed three times in PBS and counter stained with DAPI (Sigma-Aldrich) diluted to 1 µg/mL in PBS for 10 min in the dark at RT before being washed in PBS. Samples were then imaged using a BioTek Cytation 5 Multimode Reader. Wells were imaged using a 10× objective across a 7×7 montage. The images were processed and analysed using BioTek Gen5 software (version 3.08.01). Briefly, cellular analysis was performed by defining individual cells using a primary mask based on DAPI fluorescence and an appropriate object selection size (10 µm – 50 µm) and a secondary mask expanding from the DAPI-defined nuclear membrane by 30 µm. The sum intensity of COPI labelling-associated green fluorescence within the secondary mask was measured for each cell as a measure of protein abundance.

### Cell Viability Assays

Cell viability assays were performed using a CellTiter-Glo 2.0 luminescent cell viability assay (Promega) as per the manufacturer’s instructions using a BioTek Cytation 5 Multimode Reader.

### Infectivity Assays

Virus-containing cell culture supernatants were recovered from indicated experiments at the specified timepoints, clarified by centrifugation and stored at -80°C. Infectivity was assessed by focus forming assay (FFA), as previously described (REF).

### SDS-PAGE and Western blotting

Cell culture supernatants and lysates were recovered from infected and transfected Huh-7.5 cells and derivatives as detailed above. Supernatants were clarified by centrifugation (16,000 × g 5 min at 4°C), divided into aliquots, mixed with non-reducing sample buffer, boiled at 95°C for 5 min, and stored at -20°C. Lysates were harvested using NP40 lysis buffer (1% [v/v] NP-40, 50 mM Tris-HCl [pH 8.0], 150 mM NaCl) containing protease inhibitor cocktail (Sigma-Aldrich), as described previously (Eyre 2016,), divided into aliquots, mixed with reducing (for β-actin analysis) or non-reducing (for NS1 analysis) buffer, boiled at 95°C for 5 min, and stored at -20°C. Samples were separated by SDS-PAGE and transferred to nitrocellulose membranes (Bio-Rad). For cell culture supernatant normalisation, total protein stain analysis was performed using Revert™ 700 Total Protein Stain (LI-COR), following manufacturer’s instructions and imaged using a LI-COR Odyssey imaging system (Flinders Proteomics Facility, Flinders University, Australia). Following blocking (5% [w/v] skim milk in TBS for 1 h), membranes were incubated with primary antibody (anti-NS1 [1:10] or anti-β-actin [1:5000] in TBS supplemented with 0.05% [v/v] Tween-20 [Sigma-Aldrich] and 1%[w/v] skim milk) at 4°C overnight. After stringent washing, membranes were incubated with IRDye® 800 CW goat anti-mouse IgG secondary antibody (1:15,000) for 1 h in the dark at room temperature (RT). Membranes were then washed and imaged using a LI-COR Odyssey imaging system (Flinders Proteomics Facility, Flinders University, Australia), and signal intensities were quantified using Image Studio Lite (version 5.2.5).

### Immunofluorescent Labelling and Confocal Microscopy

For GCA treatment experiments, Huh-7.5 cells were infected with DENV (MOI ∼1) in a 25 cm^2^ flask. At 4 h.p.i., cells were trypsinised and re-seeded at 1 × 10^4^ cells/well into #1.5 coverglass-bottomed µ-Slide 8-well chamber slides (ibidi Gmbh, Germany) that were pre-coated with 0.2% (w/v) gelatin and returned to culture for a further 18 h. Uninfected Huh-7.5 control cells were handled similarly in parallel. At 24 h.p.i., cells were washed twice in complete DMEM and returned to culture in complete DMEM supplemented with increasing concentrations of GCA or 0.1% (v/v) DMSO vehicle control for a further 18 h. At 18 hours post-GCA treatment, cells were washed prior to fixation using ice-cold acetone:methanol (1:1) for 5 min at 4°C. Cell monolayers were then washed with PBS and blocked using 5% (w/v) BSA in PBS for 30 min at RT. Samples were labelled with primary antibodies anti-NS1 (1:5) and anti-GM130 (1:1000) diluted in 1% (w/v) BSA in PBS at 4°C overnight. Samples were then washed three times in PBS before incubation for 1 h in the dark at 4°C with AlexaFluor 488-conjugated anti-rabbit IgG and Alexa Fluor 555-conjugated anti-mouse IgG antibodies (Thermo Fisher Scientific) diluted to 1:500 and 1:1000, respectively. Samples were then washed three times in PBS and counter stained with DAPI (Sigma-Aldrich) at 1 µg/mL in PBS for 10 min in the dark at RT before being washed with PBS.

For confocal analysis of NS1 localization with respect to GFP-tagged COPA, COPB2 and COPG1, samples were prepared and labelled as above, with the exceptions that cells were seeded directly into gelatin-coated µ-Slide 8-well chamber slides, cultured overnight and infected with DENV2 (MOI ∼0.1). After culture for 24 h, cells were fixed and blocked, as above, before incubation for 1 h at RT in anti-NS1 (diluted 1:5) and anti-GFP (diluted 1:200) antibodies diluted in PBS containing 1% BSA (w/v). Cells were then washed with PBS, incubated with secondary antibodies (both diluted to 1:200) and washed, as described above. Samples were imaged using a ZEISS LSM 880 Fast Airyscan confocal fluorescence microscope system using a C-Plan-Apochromatic 63× (NA: 1.4) oil immersion objective (Flinders Microscopy and Microanalysis, Flinders University, Australia). For GCA treatment samples, laser lines 405 nm, 488 nm and 561 nm were used at 2% maximal power with appropriate detector master gain settings to enable signal visualisation with minimal saturation. Pinhole sizes were set to 1.0 Airy units for the longest-wavelength fluorophore and matched for all tracks. For confocal analysis of NS1 localization with respect to GFP-tagged COPA, COPB2 and COPG1, the 32-channel Airyscan detector was used in combination with appropriate Main Beam Splitter, Second Beam Splitter and Emission Dual Filters, the SR Airyscan Mode and appropriately reduced Master Gain settings (<50% of the dynamic range) and laser powers to satisfy Airyscan automatic alignment requirements. For this, images were acquired using 3× zoom and Z-stack intervals of 0.15 µm and processed using 3D Airyscan Processing and automatic conditions. Images were processed and analysed using ZEN Blue (version 3.2) software (ZEISS). Where indicated, colocalisation analysis was performed by measurement of Pearson’s correlation coefficients for each cell (>30 cells/group) following drawing Bezier regions of interest around each cell using ZEN Blue. Where indicated, line profiles of 15 µm were drawn through typical NS1 foci and fluorescence intensities for both channels at each point along the line were exported and graphed using GraphPad Prism version 10.2.2 software.

### Statistical Analysis and Software

With the exception of siRNA screen data analysis (see Supplemental Material), GraphPad Prism version 10.2.2 was used for statistical analyses and graphing. Unless otherwise indicated, data are means ± S.D. Details for statistical tests are provide in Figure Legends. Figures were compiled using Adobe Photoshop version 24.1.0 software.

## ACKNOWLEDGEMENTS

We extend our sincere gratitude to Amanda L. Aloia who coordinated and directed the customised membrane-trafficking and deconvolution siRNA screen assays and performed the associated data analysis. We are grateful to the following people for generously providing reagents (as detailed in Methods and Materials): Charles M Rice (Rockefeller University); Ralf Bartenschlager (University of Heidelberg); Jody Peters (University of Queensland); Roy Hall (University of Queensland); John Aaskov (Queensland University of Technology) and; Karla J. Helbig (La Trobe University). We wish to acknowledge and extend appreciation to the staff and for the services provided at the following facilities that were used during this study: Flinders Microscopy and Microanalysis (Flinders University); Cell Screen SA Facility (Flinders University); Flinders Proteomics Facility (Flinders University); Flow Cytometry Facility (Flinders University); the Australian Genome Research Facility (Adelaide, South Australia). The authors would like to acknowledge funding support from the National Health and Medical Research Council (NHMRC, Australia) (1163662) N.S.E.

